# Deletion of class II ARF causes essential tremors through Nav1.6 traffic impairment

**DOI:** 10.1101/357160

**Authors:** Nobutake Hosoi, Koji Shibasaki, Mayu Hosono, Ayumu Konno, Yo Shinoda, Hiroshi Kiyonari, Kenichi Inoue, Shinichi Aizawa, Shinichi Muramatsu, Yasuki Ishizaki, Hirokazu Hirai, Teiichi Furuichi, Tetsushi Sadakata

## Abstract

ADP-ribosylation factors (ARFs) are a family of small monomeric GTPases consisting of three classes. In the present study, we generated class II ARF-deficient mice (ARF4^+/−^/ARF5^−/−^) and found that they exhibited severe movement-associated tremors. Treatment of the mice with propranolol and gabapentin, which alleviate symptoms in patients with essential tremors, similarly reduced the amplitude of the pathologic tremors. *In vivo* electrophysiological recordings of the ARF4^+/−^/ARF5^−/−^ mice revealed that they exhibited reduced excitability of their cerebellar Purkinje cells. Immunohistochemical studies revealed that ARF4^+/−^/ARF5^−/−^ mice exhibit a severe, selective reduction of Nav1.6 proteins that are important for maintaining repetitive action potential firing in the axon initial segments (AISs) of the Purkinje cells. This decrease in Nav1.6 protein expression and the consequent tremors were alleviated by Purkinje cell-specific expression of ARF5. These results indicate that class II ARF mediates the selective trafficking of Nav1.6 to the AISs in cerebellar Purkinje cells, and suggest that the essential tremors can be ascribed to the reduced intrinsic excitability of Purkinje cells, caused by the selective decrease of Nav1.6 proteins in the AISs.

## Introduction

Essential tremor (ET) is the most frequent movement disorder, with a prevalence of approximately 4% among adults aged 40 years and older (Louis et al, 1995). In contrast to the resting tremor that is observed in Parkinson disease, ET is characterized by postural and kinetic components (Pahwa & Lyons, 2003). Growing clinical and neuro-imaging evidence has implicated cerebellar dysfunction in the pathogenesis of ET, and emerging postmortem studies have identified structural changes in the cerebellum, particularly in PCs (Benito-Leon et al, 2009; Elble & Deuschl, 2011; Kuo et al, 2011; Louis et al, 2007). The current major model that shares some features of ET uses the GABA_A_ receptor inverse agonist, harmaline, to induce a temporary tremor in animals (Wilms et al, 1999). A major limitation of this model is the development of a rapid tolerance to harmaline and a lack of response to anti-ET medication such as propranolol (Iwata et al, 1993; O’Hearn & Molliver, 1993). GABA_A_ receptor αl subunit-deficient mice are the only genetic animal tremor models subjected to pharmacologic profiling with anti-ET medications (Kralic et al, 2005). However, detailed mechanistic examinations have not been carried out so far.

ARF proteins belong to the Ras superfamily of small GTPases and regulate protein trafficking through the secretory and endocytic pathways (Donaldson & Jackson, 2011; Mizuno-Yamasaki et al, 2012). Six ARF proteins have been identified so far, and only five are expressed in humans as ARF2 has been lost (Cockcroft et al, 1994). Based on amino acid sequence homology, the six ARFs are grouped into the following three classes: class I (ARF1–3), class II (ARF4 and 5), and class III (ARF6). Although class II ARFs have been implicated in Golgi-to-ER retrograde transport and dense-core vesicle synthesis (Sadakata et al, 2010; Volpicelli-Daley et al, 2005a), little is known about their role *in vivo*.

The aim of the current study was to clarify the role of class II ARFs on normal cerebellar functioning. Both ARF4 and ARF5 are widely expressed in the mouse brain, and ARF4 knockout (KO) mice are embryonic lethal (Jain et al, 2012). To elucidate the functional roles of class II ARFs *in vivo,* we first generated an ARF4 heterozygous (+/−) mouse line and an ARF5 KO (−/−) mouse line and crossed them to produce a class II ARF-reduced mouse line, ARF4^+/−^/ARF5^−/−^ mice. Using ARF4^+/−^/ARF5^−/−^ mice, we found that class II ARFs play an important role in the selective trafficking of Nav1.6 to the AIS and that class II ARF deficiency causes a severe ET phenotype.

## Results and Discussion

### Class II ARF-deficient mice exhibit severe movement-associated tremors

ARF4 and ARF5 KO mice were generated as described in the Materials and Methods section (Figure EV1). As previously described (Jain et al, 2012), ARF4-null homozygotes (ARF4^−/−^) were embryonically lethal, whereas the heterozygotes (ARF4^+/−^) were viable, therefore ARF4^+/−^ were used for subsequent experiments. The ARF5 KO mice generated in this study is the first ARF5 null mice. Wild-type (WT), ARF5^+/−^, and ARF5^−/−^ pups were born at the expected 1:2:1 Mendelian frequency. Neither ARF4 nor ARF5 proteins were detected in the respective KO mice, and a reduced amount was present in those of heterozygous mice (Figure EV1). There were no apparent significant phenotypic abnormalities in the ARF4^+/−^ nor the ARF5^−/−^ mice. While ARF4^+/−^/ARF5^−/−^ and WT mice had a similar life expectancy, performance on the Rota-Rod revealed that the double mutants presented motor deficits (Figure 1A). In addition, the ARF4^+/−^/ARF5^−/−^ mice exhibited severe tremor during movement (but not at rest) after 3–4 weeks of age (Movie EV1).

**Figure 1.**
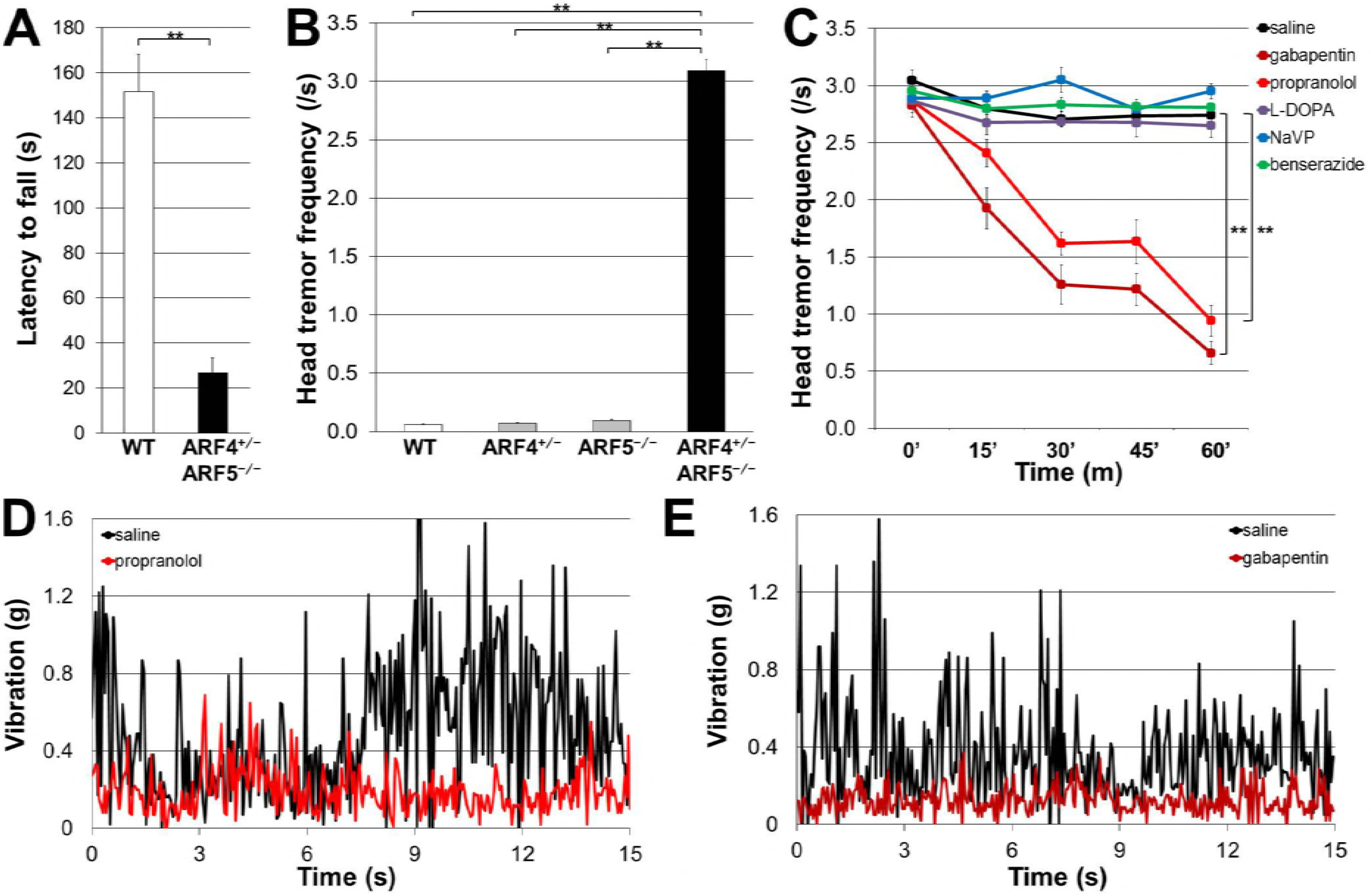
Treatment of ARF4^+/−^/ARF5^−/−^ mice with propranolol and gabapentin reduced the amplitude of the pathologic tremors. **(A)** Rotarod performance of WT (*n* = 8) and ARF4^+/−^/ARF5^−/−^ mice (*n* = 5) at 8 weeks of age (P8w). \*\**P* < 0.01, Student’s *t*-test. **(B)** Head tremor frequency of mouse during movement within 3 minutes was counted from video images. WT (white, *n* = 12), ARF4^+/−^ (gray, *n* = 12), ARF5^−/−^ (gray, *n* = 12), and ARF4^+/−^/ARF5^−/−^ mice (black, *n* = 12) of P7-8w were used. One-way ANOVA and post-hoc Scheffe test were used to determine statistical significance. \*\**P* < 0.01. Error bars indicate the SEM. **(C)** Indicated drugs were administrated to the ARF4^+/−^/ARF5^−/−^ mice (*n* = 12). \*\**P* < 0.01 compared to controls by repeated measure ANOVA. Error bars indicate the SEM. (**D, E**) Vibration before (*black*) and after (*red* in **D**, *brown* in **E**) administration of propranolol (D) and gabapentin (E). The tail of mouse was fixed with a piano wire and the vibration of piano wire was recorded by a vibration data logger.

We quantified tremor by measuring the frequency of head shaking during movement among the four genotypes (WT, ARF4^+/−^, ARF5^−/−^, and ARF4^+/−^/ARF5^−/−^). Only the ARF4^+/−^/ARF5^−/−^ mice showed head tremor during movement (Figure 1B). This abnormal behavior resembles the primary symptoms of ET patients and a subset of Parkinson’s disease patients, and these two pathologies can be separated pharmacologically. Patients with ET can be treated to reduce their tremors with several drugs, including propanolol and gabapentin (Pahwa & Lyons, 2003). Both the head shaking frequency and body vibration exhibited by ARF4^+/−^/ARF5^−/−^ mice were decreased by administration of propanolol or gabapentin (Figure 1, C-E). However, L-DOPA/benserazide (effective for improving Parkinson’s disease symptoms (Hwang et al, 2005)), sodium valproate (reported to improve cortical myoclonic tremor (Cen et al, 2016)), benserazide alone, or saline treatment had no effect on the tremors (Figure 1, C-E).

### Class II ARF-deficient mice exhibit movement-related abnormal brain activity and reduced excitability of cerebellar Purkinje cells

We next examined how the brain activity of ARF4^+/−^/ARF5^−/−^ mice was altered during movement by recording the electroencephalogram (ECoG) and electromyography (EMG) under freely moving conditions (Figure 2, A and B). We did not find any differences in the ECoG between WT and ARF4^+/−^/ARF5^−/−^ mice during non-moving (resting) periods (Figure 2, A-C). However, ECoG was dramatically changed during movement periods in ARF4^+/−^/ARF5^−/−^ mice (Figure 2, A, B and D). Analysis of ECoG power spectrums revealed a significant increase in delta-wave power in moving ARF4^+/−^/ARF5^−/−^ mice as compared to WT mice (Figure 2D). Alpha and beta-wave power were also increased during ARF4^+/−^/ARF5^−/−^ mice movement, while the theta-wave power was reduced (Figure 2D). These results indicate that basal brain activities of ARF4^+/−^/ARF5^−/−^ mice were normal (Figure 2C), but that movement triggered abnormal brain activities (Figure 2D). We also analyzed EMG power spectrums in moving WT (Figure 2E) and ARF4^+/−^/ARF5^−/−^ mice (Figure 2F) and found that EMG median power in ARF4^+/−^/ARF5^−/−^ mice was significantly different than that in WT mice (Figure 2, E and F). In our study, the electrodes for ECoG were implanted into the cerebrum, and the reference electrode into the cerebellum (Shibasaki et al, 2015). This recording configuration would mean that the ECoG in our hands reflects differential neuronal activities between the cerebrum and the cerebellum. Therefore, it is possible that the abnormality of ECoG (Figure 2, A-D) arises from the abnormal neuronal activity in the cerebellum.

**Figure 2.**
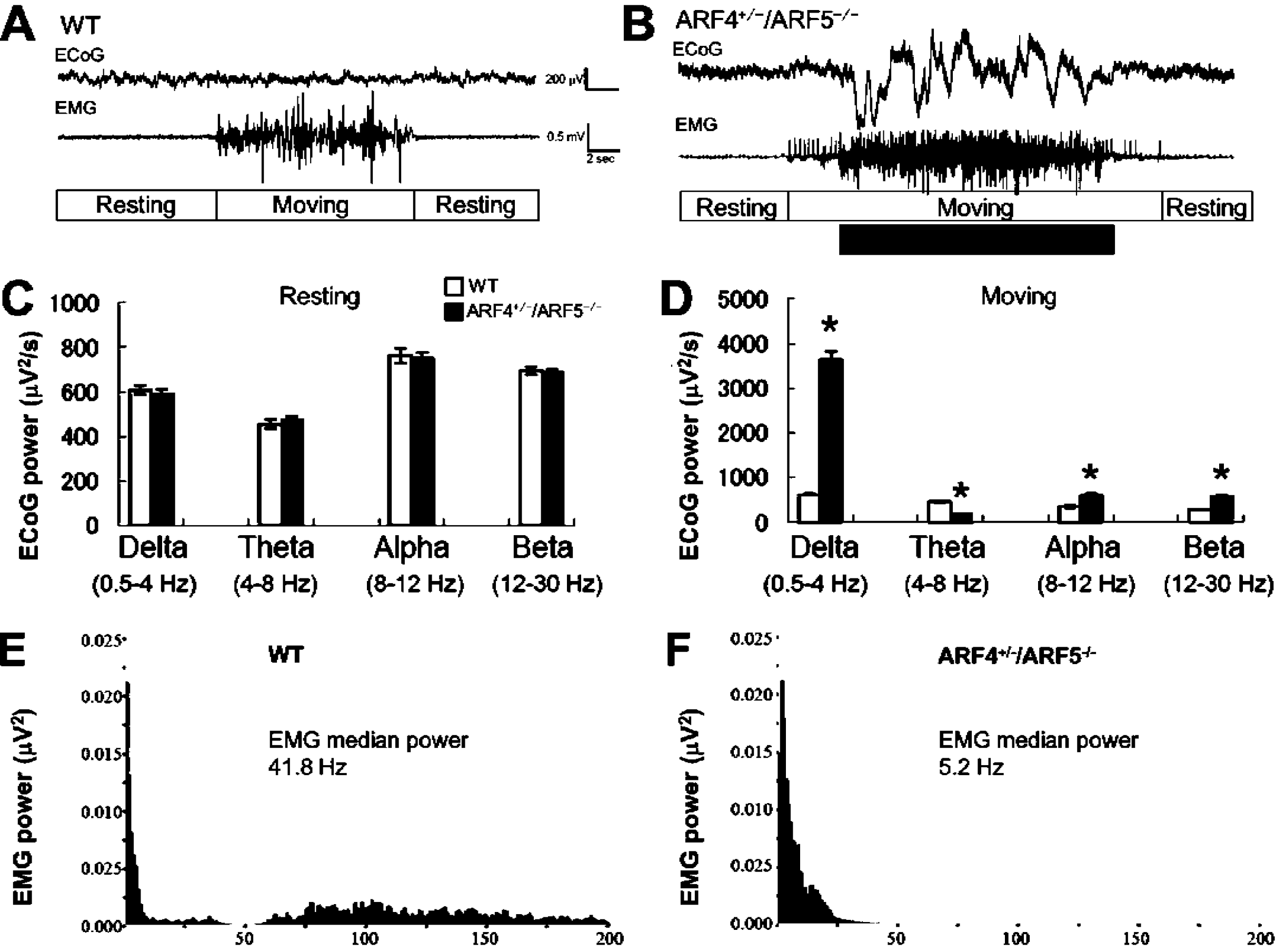
Abnormal ECoG and EMG activities in ARF4^+/−^/ARF5^−/−^ mice upon moving periods. **(A, B)** Representative ECoG and EMG traces in WT or ARF4^+/−^/ARF5^−/−^ mouse upon wakefulness. **(C, E)** ECoG power spectrums were quantified in non-moving, resting mice (Resting) and moving mice (Moving) upon wakefulness. The asterisks indicate statistically significant differences between the genotypes (WT vs. ARF4^+/−^/ARF5^−/−^, *n* = 6 each, *t*-test). **(E, F)** A representative EMG power spectrum was calculated in moving WT or ARF4^+/−^/ARF5^−/−^ mouse upon wakefulness.

Interestingly, previous studies have indicated that tremor and motor deficits may be caused by reduced intrinsic excitability of cerebellar PCs (Kalume et al, 2007; Levin et al, 2006). Thus, we recorded APs of PCs in cerebellar slices to examine this possibility in ARF4^+/−^/ARF5^−/−^ mice. Injection of depolarizing currents smaller than 600 pA evoked similar AP firing between WT and ARF4^+/-^/ARF5^−/−^ mice (Figure 3, A and B). However, injection of larger depolarizing currents resulted in much fewer spike discharges in ARF4^+/−^/ARF5^−/−^ mice (Figure 3A and B). This data indicates that the PC intrinsic excitability is indeed reduced in the case of larger current inputs in ARF4^+/−^/ARF5^−/−^ mice. This phenomena is possibly related to the behavioral phenotype in ARF4^+/−^/ARF5^−/−^ mice, where tremor-like head shaking occurs only during locomotion, but not at rest as previously descrived. Locomotion increases cerebellar granule cell activity (*i.e.*, excitatory inputs to PCs) and consequently, some PCs increase their simple spike firing rates during movement *in vivo* (Jelitai et al, 2016). Therefore, PCs are likely to receive fewer spontaneous synaptic current inputs at rest while receiving many more excitatory synaptic current inputs during locomotion. In that context, it is possible that tremors occur in ARF4^+/−^/ARF5^−/−^ mice during movement because their PCs may be unable to properly fire APs in response to movement-driven “increased” synaptic inputs.

**Figure 3.**
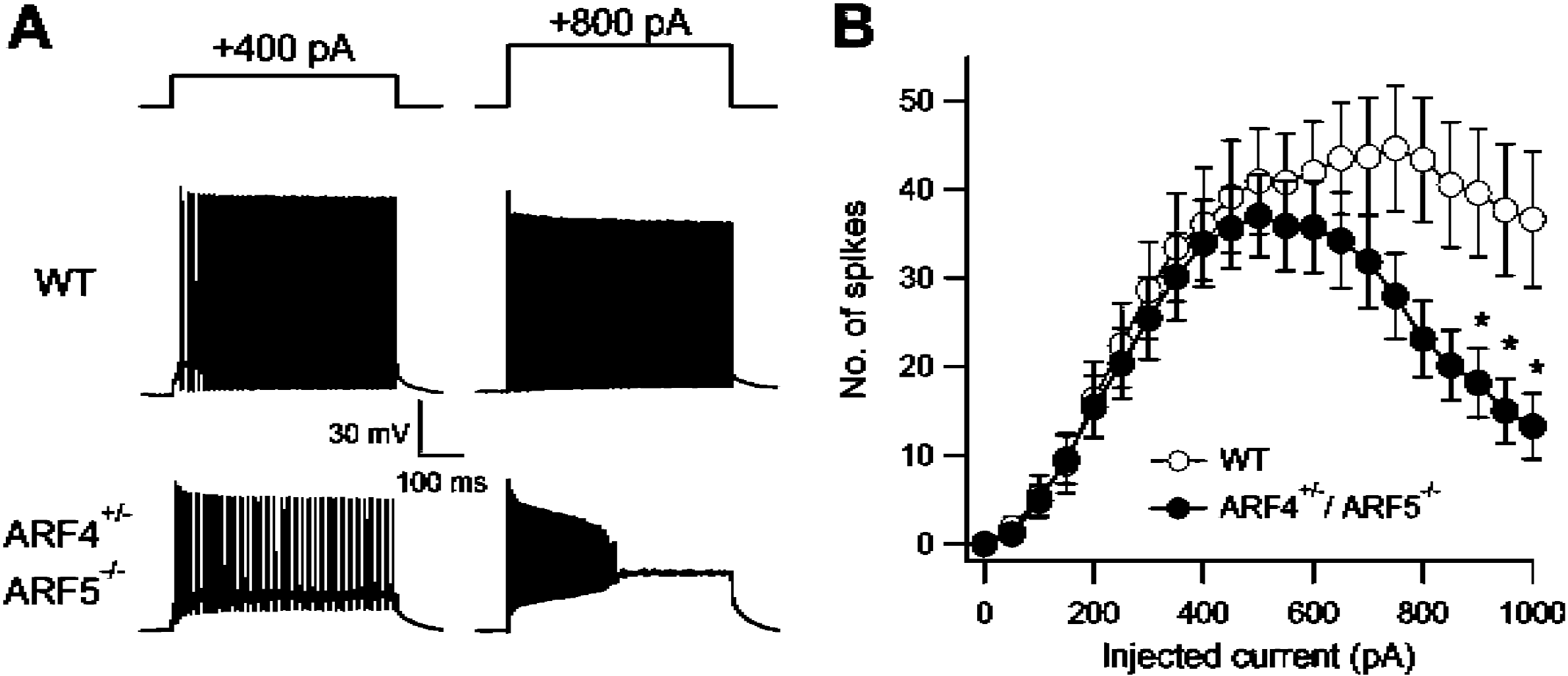
Reduced intrinsic excitability of cerebellar PCs in class II ARF-deficient mice. **(A)** Representative traces of AP firing in response to smaller (Left) and larger (Right) depolarizing current injections in WT (gray) and ARF4^+/−^/ARF5^−/−^ (black) PCs. **(B)** The average number of evoked APs plotted against the injected current amplitudes in WT (white, *n* = 27) and ARF4^+/−^/ARF5^−/−^ (black, *n* = 26) PCs. The data were collected from more than 6 mice in each condition. A repeated-measures two-way ANOVA indicates a significant interaction (genotype × injected current; F_22,1122_ = 2.671; *P* < 0.0001). Multiple comparison tests with Holm-Sidak’s method show significant differences between WT and ARF4^+/−^/ARF5^−/−^ at injected current strength indicated by *(*P* < 0.05).

We also examined the parameters of AP waveforms in WT and ARF4^+/−^/ARF5^−/−^ PCs. We found no significant differences in most of the parameters, such as maximum rate of rise, maximum rate of fall, AP threshold, AP half width, and afterhyperpolarization (minimum voltage of AP), between WT and ARF4^+/−^/ARF5^−/−^ mice, except that AP peak was smaller in ARF4^+/−^/ARF5^−/−^ mice (Table 1). The passive electrical properties of PCs, which may affect AP output (Bekkers & Hausser, 2007), were similar between WT and ARF4^+/−^/ARF5^−/−^ mice (Table 1). These results suggest that some active conductance mediated by voltage-dependent sodium or other ion channels, may be altered in ARF4^+/−^/ARF5^−/−^ PCs.

**Table 1.** Action potential (AP) parameters and passive electrical properties in WT and ARF4^+/−^/ARF5^−/−^ PCs. The first APs evoked by the smallest injected currents were analyzed to estimate AP waveform parameters above. Maximum rates of rise and fall in each AP were measured by detecting positive and negative peak values of its differentiated waveform (dV/dt), respectively. The AP threshold was defined as the membrane potential at which dV/dt exceeded 20 mV/ms. AP half amplitude width was defined as the width at the midpoint between the threshold and the peak of its AP. The passive electrical properties of PCs were estimated using averaged traces of around 20 current responses evoked by hyperpolarizing voltage pulses (from −70 to −75 mV, 500 ms duration) in a voltage clamp mode.

| **Action potential (AP) parameters** |  |  |  |
| --- | --- | --- | --- |
|  | WT (*n* = 29) | ARF4<sup>+/-</sup>/ARF<sup>-/-</sup> (*n* = 26) | *t*-test results |
| AP peak (mV) | 31.0 ± 1.4 | 26.8 ± 1.6 | *p* < 0.05 |
| Maximum rate of rise (mV/ms) | 404.9 ± 18.2 | 373.8 ± 14.0 | *p* = 0.181 |
| Maximum rate of fall (mV/ms) | -268.7 ± 16.1 | -248.0 ± 11.9 | *p* = 0.306 |
| AP threshold (mV) | -43.2 ± 1.0 | -45.2 ± 1.1 | *p* = 0.205 |
| AP half amplitude width (ms) | 0.36 ± 0.02 | 0.36 ± 0.01 | *p* = 0.900 |
| AP afterhyperpolarization (mV) | -58.0 ± 1.0 | -59.2 ± 1.1 | *p* = 0.422 |
| Resting membrane potential (mV)<br>before current steps | -66.3 ± 0.45 | -67.0 ± 0.48 | *p* = 0.288 |
| **Passive electrical properties** |  |  |  |
|  | WT (*n* = 38) | ARF4<sup>+/-</sup>/ARF<sup>-/-</sup> (*n* = 30) | *t*-test results |
| Membrane capacitance (pF) | 552.1 ± 45.6 | 512.3 ± 46.8 | *p* = 0.545 |
| Input resistance (MΩ) | 239.2 ± 32.9 | 272.3 ± 36.7 | *p* = 0.504 |

### Class II ARF-deficient mice exhibit selective depletion of Nav1.6 proteins in the axon initial segments (AISs) of Purkinje cells

Cerebellar PCs have two types of pore-forming α subunits (Na_v_1.1 and Na_v_1.6) in their voltage-gated sodium channels, which are responsible for AP initiation and propagation (Schaller and Caldwell, 2003). Similar to ARF4^+/−^/ARF5^−/−^ mice, Nav1.6-deficient mice exhibit ataxia, tremor, complete hind limb paralysis, and reduced action potential (AP) firing in PCs (Levin et al, 2006; Meisler & Kearney, 2005). We therefore performed immunohistochemical analysis of Nav1.6 expression in ARF4^+/−^/ARF5^−/−^ mice. Nav1.6 expression in the AIS was severely decreased in ARF4^+/−^/ARF5^−/−^ PCs (Figure 4, A-G). However, there was no significant difference in the density of Nav1.6 puncta in the dendrites or the soma (Figure EV2) of PCs between WT and ARF4^+/−^/ARF5^−/−^ mice. Moreover, no differences were observed between WT and ARF4^+/−^/ARF5^−/−^ Nav1.6 puncta in the white matter of their cerebella (Figure EV3). These results suggests that class II ARF mediates specifically, the trafficking of Nav1.6 to the AISs of cerebellar PCs.

**Figure 4.**
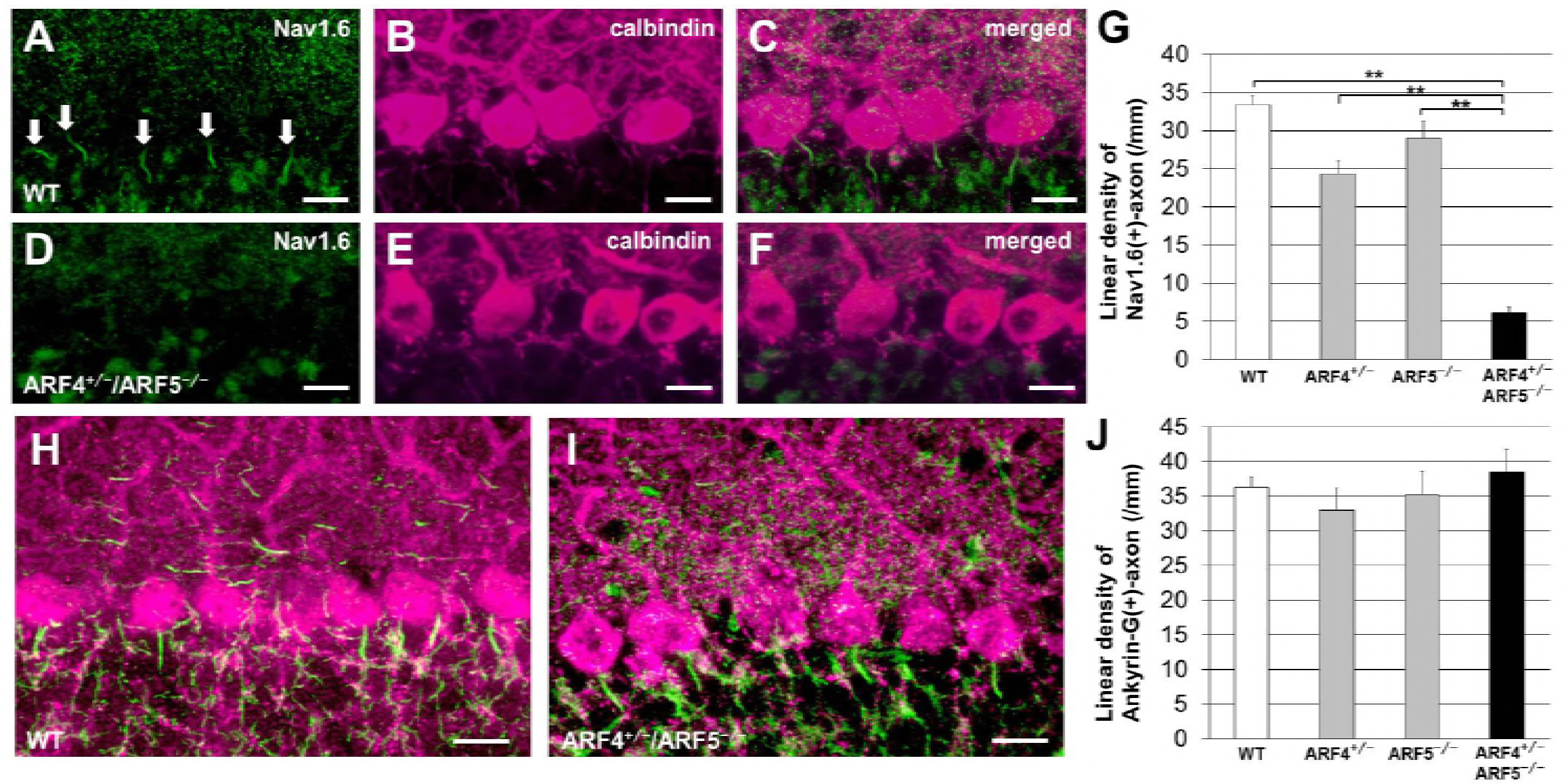
Reduced Nav1.6 localization at AIS of PCs in class II ARF-deficient mice. **(A-F)** Sagittal sections of P8w WT (A-C) and ARF4^+/−^/ARF5^−/−^ (D-F) mouse cerebella were immunolabeled with an anti-Nav1.6 antibody (*green*) and an anti-calbindin (*magenta*) antibody. White arrow indicates Nav1.6 immunoreactivities in the AIS of Purkinje cells. Scale bars, 20μm. **(G)** The linear density was determined by counting the number of Nav1.6-positive axons of Purkinje cells per millimeter line length throughout the section. WT (white, *n* = 41), ARF4^+/−^ (gray, *n* = 57), ARF5^−/−^ (gray, *n* = 36), and ARF4^+/−^/ARF5^−/−^ mice (black, *n* = 36) at P8w. One-way ANOVA, post-hoc Scheffe test, \*\**P* < 0.01. **(H-I)** Sagittal sections of P8w WT (H) and ARF4^+/−^/ARF5^−/−^ (I) mouse cerebella were immunolabeled with an anti-ankyrin-G antibody (*green*) and an anti-calbindin (*magenta*) antibody. Scale bars, 20 μm. (**J**) The linear densities of ankyrin-G-positive axon of Purkinje cells for WT (white, *n* = 13), ARF4^+/−^ (gray, *n* = 16), ARF5^−/−^ (gray, *n* = 12), and ARF4^+/−^/ARF5^−/−^ mice (black, *n* = 13) at P8w. Error bars indicate SEM.

Ankyrin-G (also known as ankyrin-3) binds to the cytoplasmic loop II-III of sodium channels to recruit and localize the channels to the AIS and the nodes of Ranvier (Dzhashiashvili et al, 2007). Surprisingly, when we examined ankyrin-G localization by immunohistochemistry, we found that ankyrin-G proteins were localized normally in the AISs of ARF4^+/−^/ARF5^−/−^ PCs (Figure 4, H-J).

We next examined the localization of other Nav subtypes in the axons of PCs using an anti-pan-Nav1 antibody. As compared to WT, similar immunoreactivities were identified at the AIS of PCs (Figure EV4). These results suggest that other Nav subtypes may compensate for the reduction of Nav1.6 at AISs in ARF4^+/−^/ARF5^−/−^ PCs.

The K_v_3 family of potassium channels is suggested to maintain high-frequency spiking in cerebellar PCs (Joho & Hurlock, 2009). In particular, Kv3.3=, is suggested to be the dominant subtype expressed in the somas of adult cerebellar PCs (Chang et al, 2007; Joho & Hurlock, 2009; McKay & Turner, 2005; Southan & Robertson, 2000). Immunohistochemical analysis revealed normal expression pattern of the Kv3.3 proteins in ARF4^+/−^/ARF5^−/−^ PCs (Figure EV5). Collectively, these immunohistochemical results indicate that impaired accumulation of Nav1.6 in AISs may underlie the reduced PC excitability in ARF4^+/−^/ARF5^−/−^ mice.

Because cerebellar PCs of Nav1.6-null mutant mice express Nav1.1 at their AISs and nodes of Ranvier in compensation for loss of the Nav1.6 subunit (Van Wart & Matthews, 2006), class II ARF-null PCs also might express compensatory Nav1.1 subunits at their AISs. If this is the case, because Nav1.1 and Nav1.6 have similar activation gating parameters (Patel et al, 2015), no differences are expected between WT and ARF4^+/−^/ARF5^−/−^ PCs regarding the AP threshold and maximum rate of rise. Our results are in line with this assumption (Table 1). The compensatory expression of Nav1.2 is unlikely, because PCs lack Nav1.2 (Kalume et al, 2007; Lorincz & Nusser, 2008) and Nav1.2 has a much higher activation threshold than Nav1.6 or Nav1.1 (Rush et al, 2005). If Nav1.2 subunits were expressed at the AIS of ARF4^+/−^/ARF5^−/−^ PCs, AP threshold in ARF4^+/−^/ARF5^−/−^ mice would have differed from that of WT mice, which was not observed in the present study. In addition, no differences in Kv3.3 (the dominant Kv3-type channels in PCs) immunoreactivity were observed between WT and ARF4^+/−^/ARF5^−/−^ PCs (Figure EV5). The AP repolarization phase (*i.e.*, AP fall) and AP afterhyperpolarization are mediated by Kv channels (Joho & Hurlock, 2009). We found no differences in AP fall and afterhyperpolarization in our study (Table 1), indicating that the Kv channels involved in AP generation are unaffected in the PCs of ARF4^+/−^/ARF5^−/−^ mice.

### Decrease in Nav1.6 protein expression and the consequent tremors alleviated by Purkinje cell-specific expression of ARF5

To examine whether the morphological and behavioral defects of ARF4^+/−^/ARF5^−/−^ mice were actually caused by the loss of class II ARF protein in cerebellar PCs, we expressed ARF5 proteins specifically in PCs of ARF4^+/−^/ARF5^−/−^ mice using adeno-associated virus serotype-9 (AAV9) expression vectors. AAV9 expressing ARF5 tagged with HA under the control of PC-specific L7 promoter, was directly administered into the ARF4^+/−^/ARF5^−/−^ cerebellum at P14 as described previously (Sawada et al, 2010). The expression of the transgene in the cerebellum was examined 6 weeks after the injection. Figure 5A and 5B shows representative sagittal sections of the cerebellar vermis from ARF4^+/−^/ARF5^−/−^ mice injected with the AAV vectors. ARF5 was diffusely expressed in all lobules of the ARF4^+/−^/ARF5^−/−^ mouse cerebellar vermis (Figure 5A) and its expression was limited to PCs (Figure 5B). The frequency of head shaking of the AAV-injected mice was significantly decreased as compared with that of the PBS-injected mice (Figure 5C). Concomitantly, the linear density of Nav1.6 expression in the AISs of PCs was increased in the ARF5-restored ARF4^+/−^/ARF5^−/−^ mice, as compared with that in the PBS-injected mice (control) (Figure 5D-F). Thus, the abnormal behavioral and immunohistochemical phenotypes (*i.e.*, tremor and Nav1.6 mislocalization) were partly rescued by PC-specific class II ARF protein expression. Therefore, we conclude that class II ARF deficiency in cerebellar PCs can contribute to ET directly through impaired localization of Nav1.6 to their AISs.

**Figure 5.**
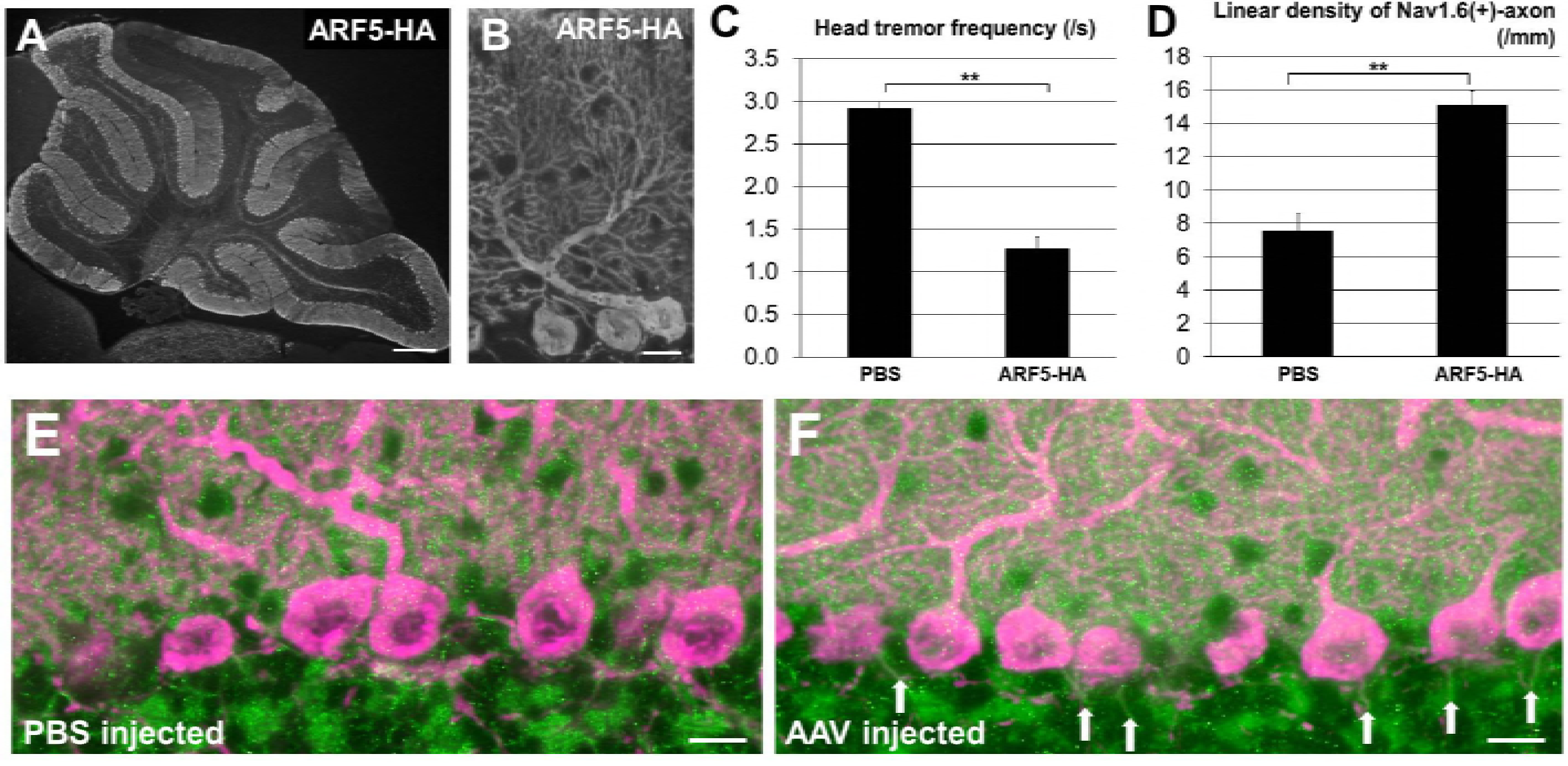
Rescue of ARF4^+/−^/ARF5^−/−^ phenotypes by exogenous ARF5 expression. **(A, B)** Sagittal section of a P8w mouse cerebellum that received injection of AAV9 vectors expressing ARF5-HA at P14 and immunolabeled with an anti-HA antibody (*green*). Scale bars are 300 μm (A) and 20 μm (B). **(C)** Head tremor frequency of P8w mouse during 3 minutes of movement was counted from video recordings. ARF4^+/−^/ARF5^−/−^ mice were injected PBS (*n* = 12) or AAV9 vectors expressing ARF5-HA (*n* = 12) at P14. \*\**P* < 0.01, Student’s *t*-test. **(D)** The linear densities of Nav1.6-positive axons of Purkinje cells for ARF4^+/−^/ARF5^−/−^ mice injected with PBS (*n* = 36) and AAV9 vectors expressing ARF5-HA (*n* = 60) at P8w. \*\**P* < 0.01, Student’s *t*-test. Error bars indicate the SEM. **(E, F)** Sagittal sections of P8w ARF4^+/−^/ARF5^−/−^ mouse cerebella injected with PBS (E) or AAV9 vectors expressing ARF5-HA (F) immunolabeled with an anti-Nav1.6 antibody (*green*) and an anti-calbindin (*magenta*) antibodies. White arrow indicates Nav1.6 immunoreactivities in the AIS of PCs. Scale bars, 20 μm.

### Function of class II ARFs

Little is known about the role of the class II ARFs in membrane trafficking. It has been documented that class II ARFs are involved in Dengue virus secretion (Kudelko et al, 2012) and hepatitis C virus replication (Farhat et al, 2016), although the specific mechanisms are unclear. In this study, we identified for the first time that class II ARF regulates selective Nav1.6 protein trafficking to the AIS (Figure 4). The AIS has two important physiological functions. One is to initiate APs, and another is to form an axonal barrier (Yoshimura & Rasband, 2014). In our analysis, class II ARF was not involved in the formation of the AIS (Figure 4), suggesting that ARF4^+/−^/ARF5^−/−^ PCs have normal axonal barrier functions to prevent nonspecific transport from the soma to axon. Moreover, our results suggest that the class II ARF is not involved in targeting of Nav1.6 to the somatodendritic compartment nor to the nodes of Ranvier (Figures EV2 and EV3). Notably, our data clearly indicates that class II ARFs specifically regulate Nav1.6 targeting to the AIS. Therefore, class II ARFs may play a role as ‘ion channel organizers’ at the AIS. It was previously suggested that AIS-localized Nav protein is first inserted in the somatodendritic compartments and subsequently endocytosed and restricted to the AIS (Fache et al, 2004). On the other hand, ARF4 and ARF5 double knockdown displayed impaired endosome recycling into the plasma membranes *in vitro* (Volpicelli-Daley et al, 2005b). Taken together, these results suggest that class II ARFs mediate the fusion of endosomes containing Nav1.6 proteins to the AIS membrane of cerebellar Purkinje cells, although the underlying mechanism remains to be studied in a greater detail.

### Essential tremor and cerebellar Purkinje cells

Kralic et al. (2005) reported a similar ET-like whole body tremor in GABA_A_ receptor α1 (*Gabra1*) subunit-deficient mice, which displayed a loss of inhibitory responses in cerebellar PCs. A decisive cause and a critical brain area (or cell type) involved in the etiology of ET were not defined in *Gabra1* KO mice probably because a global KO was used without a genetic rescue experiment (Kralic et al, 2005). Since *Gabra1* KO mice exhibited no abnormalities in the density, gross morphology, or spike firing of the PCs, their ET-like tremor might result from the impairment of GABAergic inhibition somewhere in the motor pathway and not in cerebellar PCs (Kralic et al, 2005). On the other hand, making use of the AAV-mediated PC-specific gene-rescue experiment, we could demonstrate the contribution of cerebellar PCs to the tremor pathology in class II ARF-deficient mice (Figure 5). However, the PC-specific genetic rescue did not fully correct the tremor phenotype in ARF-deficient mice, probably because not enough class II ARF proteins are expressed in the whole cerebellum. Alternatively, a possible abnormality in other regions, including the motor pathway, in addition to cerebellar PCs may be involved in the ET pathology, as proposed by Kralic et al. (2005). The abnormal ECoG observed in class II ARF-deficient mice may reflect the aberrant AP firing of neurons in the cerebral cortex to which we did not deliver the rescue vector virus in the present study.

Finally, it should be noted that the pathologic tremor of our class II ARF-deficient mice showed a remarkable pharmacological similarity with that of human patients with ET. Thus, the dysfunction of class II ARFs and the consequent disorganization of the ion channel composition (*i.e.* loss of Nav1.6) at the AIS may be few of the causes underlying human ET. We propose ARF4^+/−^/AFR5^−/−^ mice as a suitable animal model to elucidate the mechanisms of ET in humans, which may contribute to the development of future interventions.

## Materials and Methods

### Animals

All procedures for the care and treatment of animals were carried out according to the Japanese Act on the Welfare and Management of Animals and the Guidelines for the Proper Conduct of Animal Experiments issued by the Science Council of Japan. All experimental protocols were reviewed and approved by the Gunma University Animal Care and Experimentation Committee and by the Institutional Animal Care and Use Committee of the RIKEN Kobe Branch.

### Generation of ARF4 and ARF5 knockout mice and genotyping

ARF4 and ARF5 knockout (KO) mice (Accession No. CDB0884K (ARF4) and CDB0885K (ARF5); http://www2.clst.riken.jp/arg/mutant%20mice%20list.html) were generated as described (http://www2.clst.riken.jp/arg/methods.html). The targeting vector contained two loxP sites and a neomycin resistance cassette for selection in embryonic stem (ES) cells derived from C57BL/6 mice (Kiyonari et al, 2010). For gene targeting, ES cell screening and chimera production were performed. The chimeric mice were mated with C57BL/6 mice to generate F1 heterozygotes, which were then crossed with CAG-Cre mice (Sakai & Miyazaki, 1997), to produce ARF4 and ARF5 KO offspring (Figure EV1). Genomic DNA was isolated from mouse ear punched tissues and yolk sac for genotyping. ARF4 KO mouse alleles were genotyped by allele-specific PCR with the following primers: ARF4 (forward primer) 5‘-ctatgcagatggtgttaggc-3’, ARF4 (WT primer) 5‘-gccctccacatcaacacttc-3’, and ARF4 (KO primer) 5‘-caatgctaggacaatgaggc-3’. The ARF4 (forward) and ARF4 (WT) primer set yields a product of 140 bp, while the ARF4 (forward) and ARF4 (KO) primer set yields a product of 556 bp. ARF5 KO mouse alleles were genotyped using allele-specific PCR with the following primers: ARF5 (forward primer) 5‘-acttgagaaatgggtcaccg-3’, ARF5 (WT primer) 5‘- cgatctctgcaaaggacaca-3’, and ARF5 (KO primer) 5‘-cgagggaaaagctgtgttgt-3’. The primer set ARF5 (forward) and ARF5 (WT) yields a product of 156 bp and primer set ARF5 (forward), ARF5 (KO) yields a product of 280 bp. All of the engineered animals studied were backcrossed onto C57BL/6J for >5 generations.

### Antibodies

Rabbit polyclonal anti-ARF4 and anti-ARF5 antibodies were raised against mouse ARF4 (ERIQEGAAVLQKMLLEDC) and ARF5 (ERVQESADELQKMLQEDC) peptide-KLH conjugates, respectively. These were then affinity-purified against MBP-tagged full-length ARF4 and ARF5 proteins covalently coupled to CNBr-activated Sepharose 4B, and were subsequently used for Western blotting (1:1,000). The following primary antibodies were used for immunohistochemistry: mouse monoclonal anti-calbindin (1:500; cat. no. 214011, Synaptic Systems, Göttingen, Germany); rabbit polyclonal anti-Nav1.6 (1:400; cat. no. P3088, Sigma-Aldrich, St. Louis, MO); mouse monoclonal pan-Nav1 (1:4; cat. no. 73-405, Antibodies Inc., Davis, CA); mouse monoclonal anti-Kv3.3 (1:300; cat. no. 75-354, Antibodies Inc.); mouse monoclonal anti-ankyrin-G (1:300; cat. no. 75-147, Antibodies Inc.); rabbit anti-ankyrin-G (1:500; cat. no. 386003, Synaptic Systems); and rat monoclonal anti-HA (1:250; cat. no. 1867423; Roche, Mannheim, Germany).

### Electroencephalogram (EEG) and electromyogram (EMG) recording and analysis

EEG electrodes that were connected to a 5 mm^2^ computer circuit board were implanted into the cortex, and the reference electrode into the cerebellum (Shibasaki et al, 2015). In addition, electrodes for EMG were implanted into the neck muscle. The mice were then housed separately for a recovery period of at least 7 days. After 7 days, the mice were connected to an EEG machine (EEG4214, Nihon Koden, Japan), and cortical EEG (ECoG) and EMG were recorded under freely moving conditions. The ECoG and EMG recordings were taken and matched chronologically with behaviors obtained by infrared cameras. The outputs of the ECoG and EMG were connected to a PowerLab8/30 (AD Instruments, Colorado Springs, CO) to digitally convert the analog data. The digital EEG data were analyzed using the LabChart 8.0 software. The data was categorized into either stationary or moving conditions according to the EMG and behavioral movies. The EMG power spectrums were then calculated using the Labchart8.0 software.

### Cerebellar slice electrophysiology

Parasagittal slices (250–300 μm in thickness) of the cerebellar vermis were prepared from adult mice (7–20 weeks old) as described previously (Mitsumura et al, 2011), except that cutting solution (pH adjusted to 7.4 with HCl) contained NMDG (93 mM), KCl (2.5 mM), CaCl_2_ (0.5 mM), MgCl_2_ (10 mM), NaH_2_PO_4_ (1.25 mM), NaHCO_3_ (30 mM), HEPES (20 mM), and D-glucose (20 mM). The cerebellar slices were incubated at 32 °C for 20-30 min in the above solution supplemented with sodium ascorbate (5 mM), thiourea (2 mM), and sodium pyruvate (3 mM) (Zhao et al, 2011), and bubbled with 95% O_2_ and 5% CO_2_. The slices were then incubated at room temperature for more than 1 hour before recording in artificial cerebrospinal fluid (ASCF, pH 7.4) consisting of NaCl (125 mM), KCl (2.5 mM), CaCl_2_ (2 mM), MgCl_2_ (1 mM), NaH_2_PO_4_ (1.25 mM), NaHCO_3_ (26 mM), D-glucose (10 mM), bubbled with 95% O_2_ and 5% CO2, and supplemented with L-ascorbic acid (0.4 mM), myoinositol (2 mM), and sodium pyruvate (2 mM). Whole-cell recordings were performed at room temperature from the somata of Purkinje cells (PCs) using patch pipettes (2–4 MΩ) pulled from borosilicate glass (Harvard Apparatus, Holliston, MA) or #0010 glass (World Precision Instruments, Sarasota, FL). The pipette solution (pH 7.3) contained potassium gluconate (135 mM), HEPES (10 mM), KCl (5 mM), NaCl (5 mM), Mg-ATP (5 mM), Na-GTP (0.5 mM), EGTA (0.1 mM), and 0–5 phosphocreatine. During recordings, picrotoxin (100 μM) was added to the ACSF extracellular solution to block inhibitory synaptic transmission mediated by GABA_A_ receptors. Membrane potential was recorded from the patched PCs in the current-clamp mode of a MultiClamp 700B amplifier (Molecular Devices, Sunnyvale, CA) with its bridge balance and capacitance neutralization circuit. Reported membrane potentials were not corrected for junction potential (−14.6 mV). Action potentials (APs) were evoked by injecting depolarizing current steps from 0 to 1000 pA in 50 pA increments every 4–8 s. PC membrane potentials before the current injection were held at −66 mV in order to minimize voltage-dependent changes in the availability of ion channels. AP occurrences were detected by a level-crossing threshold of −10 mV. All the electrical signals were low-pass filtered at 10 kHz and sampled at 50 kHz with Digidata 1440A and pCLAMP10 software (Molecular Devices). Data analysis was performed using pCLAMP10 and Igor Pro (Wavemetrics, Inc., Lake Oswego, OR) with Neuromatic software (http://www.neuromatic.thinkrandom.com/) and custom-written Igor procedures.

### Behavioral tests

Mice were housed with a 12:12 hr light-dark cycle, with the dark cycle occurring from 20:00 to 8:00. All mice used in the experiments below were littermates from mated heterozygotes unless otherwise noted. The experimenter was blind to the genotype in all behavioral tests.

The head shake monitoring was performed as previously described (Dursun & Handley, 1993). Mice that were eight to ten weeks old were habituated to the observation cages for 60 min before video image recording. Test mice received saline, propranolol (10 mg/kg, i.p.), gabapentin (40 mg/kg, i.p.), benserazide (12.5 mg/kg, i.p.), L-DOPA (25 mg/kg, i.p.), and sodium valproate (300 mg/kg, i.p.). L-DOPA was administrated 20 min after benserazide injection. Head shakes during movement were counted from video recordings played at 2x slow motion. The tails of mice were fixed with piano wire, and the vibration of the piano wire was recorded by a 3-axis vibration data logger (cat. no. DT-178A, Sato Shouji Inc., Tokyo, Japan).

Coordination and motor skills of the mice were assessed by an accelerated rotarod test. The Rota-Rod Treadmill (Muromachi Kikai, Tokyo, Japan) consisted of a gridded plastic rod (3 cm in diameter, 10 cm long) flanked by two large round plates (50 cm in diameter). The rod accelerated from 0 to 40 revolutions per minute for 3 min and remained at the top speed for 1 additional minute. Each test consisted of 4 trials with a 10 min rest between each trial, and the time that each mouse spent on the rod was recorded.

### Immunohistochemistry

C57BL/6J male mice were deeply anesthetized with an overdose of diethyl ether and transcardially perfused with phosphate-buffered saline (PBS) and then with Zamboni’s fixative (2% paraformaldehyde in 0.1 M phosphate buffer, pH 7.4, containing 0.2% picric acid). Tissues were dissected, post-fixed in Zamboni’s fixative at 4 °C for 5 h, and cryoprotected by immersion in 15% sucrose in PBS overnight at 4 °C. After embedding in Tissue-Tek OCT compound (Sakura Finetek, Tokyo, Japan), tissues were frozen on dry ice powder, and sectioned at a thickness of 14 μm using a cryostat (CM1950, Leica Microsystems, Frankfurt, Germany) held at −18 °C. The sections were air-dried for 1 h and rinsed in PBS three times. After blocking with 5% bovine serum albumin (BSA) and 0.3% Triton X-100 in PBS at room temperature for 1 h, the sections were incubated at 4°C overnight with the primary antibodies in immunoreaction buffer (2× PBS containing 0.3% Triton X-100 and 1% BSA). The sections were then washed in PBS, incubated at room temperature for 1 h with the appropriate secondary antibodies in immunoreaction buffer, and washed again in PBS. Stained sections were mounted in Vectashield mounting medium (Vector Laboratories, Peterborough, England) and observed under a fluorescence microscope (BX51, Olympus, Tokyo, Japan) equipped with a CCD camera (VB-7000, Keyence, Osaka, Japan). Digital images were processed using Adobe Photoshop CS5.1 software (Adobe, San Jose, CA).

### Production of AAV Vectors

To express the ARF5 gene with a C-terminal hemagglutinin (HA) tag in PCs, we constructed the adeno-associated virus (AAV) vector plasmid (pAAV-L7-4-minCMV-ARF5-HA), which was designed to express ARF5 under the control of a truncated L7 promoter with a minimal cytomegalovirus sequence (minCMV) (Sawada et al, 2010). Recombinant AAV serotype 9 (AAV9) particles were generated by the cotransfection of the following three plasmids in HEK293T cells: pAAV-L7-4-minCMV-ARF5-HA, pHelper (Stratagene, La Jolla, CA), and pAAV2/9 (kindly provided by Dr. J. Wilson). The viral particles were purified by ammonium sulfate precipitation and iodixanol continuous gradient centrifugation as described previously (Miyake et al, 2012). The genomic titer of the purified AAV9 viral particles as determined by real-time PCR was 2.32 × 10^13^ vector genomes/ml.

### Statistical analyses

Statistical analyses were performed using the Excel Statistics (Statcel 3; Social Survey Research Information Co, Ltd, Tokyo, Japan) and GraphPad Prism7 software (Graphpad Software, La Jolla, CA, USA). All data are presented as mean ± SEM. Differences between groups were analyzed using either unpaired Student’s *t*-test, one-way ANOVA followed by Scheffe post hoc test, or two-way ANOVA followed by a Holm-Sidak’s post hoc test, according to each experimental design. *P* < 0.05 was considered statistically significant.

## Acknowledgements

This study was supported by Grants-in-Aid for Scientific Research from Japan Intractable Diseases Research Foundation, Takeda Science Foundation, Sumitomo Foundation, SENSHIN Medical Research Foundation, Novartis Pharma, Life Science Foundation of Japan, Kawano Masanori Memorial Public Interest Incorporated Foundation for Promotion of Pediatrics, MEXT/JSPS KAKENHI Grant Numbers JP25110707, JP25430061, JP15H05934.

## Author contributions

NH, KS, and TS designed the experiments. NH, KS, MH, AK, YS, HK, KI, SM, and TS performed the experiments. NH, KS, and TS analyzed the data. NH, KS, and TS wrote the paper. NH, KS, SA, SM, YI, HH, TF, and TS contributed to the discussion.

## Conflict of interest

The authors have declared that no conflict of interest exists.

## Expanded View Figure legends

**Figure S1. Generation of ARF4^−/−^ and ARF5^−/−^ mice. (A)** Maps of the murine ARF4 gene, both the targeted allele and the ARF4 KO allele are shown. **(B)** Maps of the murine ARF5 gene, both the targeted allele and the ARF5 KO allele are shown. Exons are shown as white boxes. The *loxP* and *frt* sites are depicted as black and white arrowheads, respectively. **(C-F)** Immunoblot analysis of E13.5 whole body (ARF4) and P8w cerebellum (ARF5) of WT (+/+), heterozygous (+/−), and homozygous (−/−) mice. Protein lysates were immunoblotted with anti-ARF4, anti-ARF5, and anti-actin antibody.

**Figure S2. No difference in Nav1.6 somatodendritic localization in ARF4^+/−^/ARF5^−/−^ Purkinje cells. (A-B)** Sagittal sections of P8w WT (A) and ARF4^+/−^/ARF5 ^−/−^ (B) mouse cerebella were immunolabeled with an anti-Nav1.6 antibody (*green*) and an anti-calbindin (*magenta*) antibody. Scale bars, 20 μm. **(C-D)** Densities of Nav1.6-positive puncta in primary and secondary dendrites (C) and soma (D) of WT (white, dendrite; *n* = 40, soma; *n* = 48) and ARF4^+/−^/ARF5^−/−^ Purkinje cells (black, dendrite; *n* = 40, soma; *n* = 61).

**Figure S3. No difference in densities of Nav1.6 puncta in white matter. (A-B)** Sagittal sections of P8w WT (A) and ARF4^+/−^/ARF5^−/−^ (B) cerebellar white matter were immunolabeled with an anti-Nav1.6 antibody (*green*) and an anti-calbindin (*magenta*) antibody. Enlarged images are shown in insets. Scale bars, 10 μm. **(C)** Densities of Nav1.6-positive puncta in cerebellar white matter of WT (white, *n* = 34) and ARF4^+/−^/ARF5^−/−^ PCs (black, *n* = 21).

**Figure S4. Immunoreactivities of anti-pan-Nav1 antibody in the AISs of PCs. (A-F)** Sagittal sections of P8w WT (A-C) and ARF4^+/−^/ARF5^−/−^ (D-F) mouse cerebella were immunolabeled with an anti-pan-Nav1 antibody (*green*) and an anti-ankyrin-G (*magenta*) antibody. White arrow indicates pan-Nav1 immunoreactivities in the AISs of PCs. Scale bars, 20 μm.

**Figure S5. Kv3.3 immunoreactivity in WT and ARF4^+/−^/ARF5^−/−^ cerebellum. (A, B)** Sagittal sections of P8w WT (A) and ARF4^+/−^/ARF5^−/−^ (B) mouse cerebella were immunolabeled with an anti-Kv3.3 antibody. Scale bars, 10 mm.

